# Mycolactone enhances the Ca^2+^ leakage from endoplasmic reticulum by trapping Sec61 translocons in a Ca^2+^ permeable state

**DOI:** 10.1101/2021.05.12.443793

**Authors:** Pratiti Bhadra, Scott Dos Santos, Igor Gamayun, Tillman Pick, Joy Ogbechi, Belinda S. Hall, Richard Zimmermann, Volkhard Helms, Rachel E. Simmonds, Adolfo Cavalié

## Abstract

The *Mycobacterium ulcerans* exotoxin, mycolactone, is an inhibitor of co-translational translocation via the Sec61 complex. Mycolactone has previously been shown to bind to, and alter the structure of, the major translocon subunit Sec61α, and change its interaction with ribosome nascent chain complexes. In addition to its function in protein translocation into the ER, Sec61 also plays a key role in cellular Ca^2+^ homeostasis, acting as a leak channel between the endoplasmic reticulum (ER) and cytosol. Here, we have analysed the effect of mycolactone on cytosolic and ER Ca^2+^ levels using compartment-specific sensors. We also used molecular docking analysis to explore potential interaction sites for mycolactone on translocons in various states. These results show that mycolactone enhances the leak of Ca^2+^ ions via the Sec61 translocon, resulting in a slow but substantial depletion of ER Ca^2+^. This leak was dependent on mycolactone binding to Sec61α because resistance mutations in this protein completely ablated the increase. Molecular docking supports the existence of a mycolactone-binding transient *inhibited* state preceding translocation and suggests mycolactone may also bind Sec61α in its *idle* state. We propose that delayed ribosomal release after translation termination and/or translocon “breathing” during rapid transitions between the *idle* and *intermediate-inhibited* states allow for transient Ca^2+^ leak, and mycolactone’s stabilisation of the latter underpins the phenotype observed.

## Introduction

Mycolactone is the exotoxin produced by *Mycobacterium ulcerans* (*M. ulcerans*) that is central in the aetiology of Buruli ulcer (BU), a chronic necrotizing skin infection [1-3]. Most pathogenic strains isolated from BU patients living in West Africa, where the disease is most prevalent, produce mycolactone A/B. Other congeners also exist, produced by pathogenic strains found in Australia (mycolactone C) and so-called “ancestral” strains. Mycolactone A/B is the most toxic *in vitro* and is a polyketide composed of an invariant 12-membered lactone ring to which two polyketide-derived, unsaturated acyl side chains are attached [1,3]. The shorter “Northern” chain is invariant, while the longer “Southern” Chain gives rise to the various mycolactone congeners. Like all mycolactones, mycolactone A/B is a 3:2 rapidly-equilibrating mixture of isomers around the second double bond in the Southern fatty acid tails [4]. Structure-activity studies have proposed that mycolactone B is the active isoform [5].

Infection with *M. ulcerans* initially produces a painless swelling or nodule. The immunosuppressive properties of mycolactone enable the disease to progress without inflammation and fever, leading eventually to development of the necrotic lesions in cutaneous and subcutaneous tissues that are characteristic of BU [6]. Many reports support mycolactone as driving the pathogenic sequelae of BU that comprise skin ulceration, coagulative necrosis, local hypoesthesia and suppression of immune responses. Furthermore, molecular targets for mycolactone have been identified and proposed to be associated with its hallmarks [2,3]. Binding and activation of type 2 angiotensin receptors II initiates a signalling cascade that culminates in the activation of TWIK-related arachidonic acid-stimulated K^+^ channel in neurons [7]. The resulting hyperpolarisation of neurons was proposed to mediate the analgesic effects of mycolactone. In cell free assays, mycolactone binds to the Wiskott-Aldrich syndrome protein (WASP) and neural WASP (N-WASP), which are scaffold proteins involved in the remodelling of the actin cytoskeleton [8,9]. By interacting with WASP/N-WASP, mycolactone likely initiates an uncontrolled assembly of actin and defective cell-matrix adhesion, which may compromise skin integrity. Notably, the down regulation of multiple mediators of inflammation correlates solely with the continuous presence of mycolactone even after completion of antibiotic therapy against *M. ulcerans*, providing a link between mycolactone and immunosuppressive effects [10].

Subsequently, investigations using cell-free protein synthesis showed that mycolactone inhibits the Sec61-dependent protein translocation into the endoplasmic reticulum (ER) [11]. Furthermore, the blockade of Sec61-dependent translocation by mycolactone has been linked to inhibition of T cell activation and antigen presentation, providing a molecular basis for the mycolactone-induced immunosuppression [12,13]. The effects of mycolactone on ER function are however much broader. For instance, it has been shown that mycolactone down-regulates luminal Hsp70-type chaperone binding-immunoglobulin protein (BiP) in dendritic cells [14]. Furthermore, inhibition of Sec61-mediated translocation by mycolactone induces an integrated stress response via translational activation of activating transcription factor 4 [15]. Thus, a consequence of the blockade of Sec61 translocons is a stress response that eventually induces apoptosis and underlies cytotoxic effects of mycolactone. A forward genetic screen in HCT116 cells [15,16] identified mutations at a number of sites in Sec61α (D60, R66, S71, G80, S72 and Q127) that confer resistance to mycolactone-induced toxicity. Crucially, out of 33 resistant clones tested, all contained a mutation in at least one of these sites, suggesting that Sec61α is the prime target of mycolactone. Several of these mutations have been studied in detail and have been shown to reverse biological effects of mycolactone [12,15-17].

Sec61α plays a central role in the translocation of nascent polypeptides emerging from the translating ribosome. To accomplish translocation, the Sec61 complex undergoes substantial conformational changes. McKenna et al. (2016) provided the first biochemical evidence that the binding of mycolactone to Sec61α also induces conformational changes of the pore geometry [18]. Several crystal and cryo-electron microscopy structures have revealed the conformation of the native ribosome-Sec61 complex in the non-translocating (*idle*) state [19-21]. Here, the lateral gate of the Sec61α is closed and the so-called ‘plug’ helix of Sec61α seals the channel toward the lumenal side [22]. During translocation, the conformation of ribosome-Sec61 complexes is significantly different. Here, the plug is displaced to enable translocation and the lateral gate remains open for membrane access of the signal peptide or transmembrane domains of the nascent precursor polypeptide [23]. Recently, Gérard et al. [16] determined the structure of mammalian ribosome-Sec61 complexes inhibited by mycolactone via electron cryo-microscopy, where the Sec61α channel is trapped in a putative *intermediate* state. In this state, Sec61α is in a conformation where the cytosolic side of the lateral gate is open, whereas the plug still occupies the channel pore in the lumenal side. Notably this conformation closely resembles that of yeast Sec61 translocons bound to Sec62 and Sec63 [24,25], suggesting that this pose is adopted during native translocation at least in lower eukaryotes. The structure provides novel insight into the molecular mechanism of mycolactone as an inhibitor. Notably, mycolactone binds with weaker affinity to translocons from cells carrying resistance mutations in Sec61α [16]. Here, molecular dynamics simulations suggested that the single amino acid mutations modulate the conformational dynamics of Sec61α and thereby disfavour the mycolactone-bound conformation.

It is striking that the molecular targets of mycolactone have been associated with key functions in the Ca^2+^ homeostasis. For instance, Sec61α translocons behave as Ca^2+^ permeable channels that support a Ca^2+^ leakage from ER that is regulated by BiP [26]. Mycolactone may also interact with WASP/N-WASP leading to actin assembly [9], potentially impacting Ca^2+^ homeostasis because actin polymerisation in the cell periphery prevents the coupling between the ER and Ca^2+^-release activated Ca^2+^ channels, reducing the Ca^2+^ entry into the cell [27]. On the other hand, actin polymerisation enhances the Ca^2+^ mobilisation from ER that is induced by the sarco/endoplasmic reticulum Ca^2+^-ATPase (SERCA)-inhibitor thapsigargin (TG) [28]. Finally, as a lipophilic substance, mycolactone interacts with phospholipids and hence may perturb cholesterol-rich membranes more generally [29-31].

Thus, while it can be hypothesised that mycolactone might distort Ca^2+^ homeostasis in the cell, until now this area has been little explored. An early study reported a dose-dependent increase in cytosolic Ca^2+^ levels in L929 cells exposed to mycolactone [32]. Later, it was shown that mycolactone-mediated hyperactivation of Lck in Jurkat cells resulted in depletion of intracellular Ca^2+^ stores and T cell receptor downregulation [33]. As in the study with L929 cells, increased cytosolic Ca^2+^ levels were also found in resting T cells exposed to mycolactone. Additionally, mycolactone reduced the store-operated Ca^2+^ entry and the Ca^2+^ mobilisation induced by the non-selective iononophore ionomycin (IONO) in these T cells [33]. In the present study, we focused on the effects of mycolactone on Ca^2+^ leakage from the ER. Using mycolactone-resistant mutants of Sec61α [15-17], we show in Ca^2+^ imaging experiments that mycolactone specifically enhances the Sec61-mediated Ca^2+^ leakage from ER. By docking analysis, we further explore potential interaction sites for mycolactone with Sec61 translocons at the molecular level. These analyses support the mechanistic model in which mycolactone stabilises the *intermediate* (mycolactone-bound/mycolactone-inhibited) conformation of the ribosome-translocon complex, which in turn generates a Ca^2+^ leakage from the ER.

## Materials and Methods

### Cell lines

The experiments were done with a HEK-293 cell line that stably expresses the FRET-based D1ER sensor in the ER lumen (D1ER-HEK) as well as with HCT116 cells that express Sec61α mutants (D60G, R66K, S71F, S82Y, Q127K) [15,16,34]. D1ER was kindly provided by R. Y. Tsien [35] and stably expressed in HEK-293 cells to obtain the D1ER-HEK cell line [34]. D1ER-HEK cells were cultured under selection with G418 (Minimal Essential Medium, MEM (Gibco), 10% FBS, 0.5 mg/ml G418). For Ca^2+^ imaging experiments, cells were transferred to poly-L-lysine coated cover slips and maintained in culture for 2-3 days. Wild type (WT) HCT116 cells and those expressing Sec61 mutants were maintained in McCoyS 5A Medium (Gibco) and 10% FBS. HCT116 cells expressing Sec61 mutants (D60G, R66K, S71F, S82Y, Q127K) were generated by random mutagenesis with ENS in the DNA repair defective HCT116 cell line [15]. Prior to Ca^2+^ imaging experiments, HCT116 cells were plated on poly-L-lysine coated cover slips and transfected with ER calcium sensor ER-GCaMP6-150 using FuGENE HD (Promega Corp.). Experiments were carried out 1-2 days after transfection. ER-GCaMP6-150 was kindly provide by J. de Juan-Sanz and T. A. Ryan [36]. The murine macrophage cell line RAW264.7 was routinely cultured in high glucose Dulbecco’s Modified Eagle’s Medium (DMEM) supplemented with 10% FBS [11]. Cell culture was carried at 37°C in a humidified environment with 5% CO_2_.

### Reagents and cell treatment

Synthetic mycolactone (MYC) A/B was kindly provided by Y. Kishi (Harvard University) [37]. Aliquots containing 0.5 mg/ml DMSO were maintained at −80°C in the dark. Before the experiments, the mycolactone stock solution was diluted directly in complete culture medium or recording solution to the desired concentrations. Thapsigargin (TG, Thermo Fisher Scientific) and ionomycin (IONO, Thermo Fisher Scientific) were dissolved in DMSO to obtain 1 mM and 10 mM stocks, respectively. TG and IONO stock were maintained at −20°C in the dark and dilutions to the desired concentrations were made directly in the recording solution just before experiments.

FURA-2 AM (Thermo Fisher Scientific) was dissolved in DMSO to obtain a 1 mM stock solution, which was subsequently dissolved to 4 µM in a Ca^2+^ containing solution (140 mM NaCl, 5 mM KCl, 1 mM MgCl_2_, 1 mM CaCl_2_, 10 mM glucose, 25 mM HEPES, pH 7.2). Cells were loaded with FURA-2 prior imaging experiments by incubation with the solution containing 4 µM FURA-2 AM for 20 min, washed and subsequent exposure to the bath solution.

TG and IONO were routinely applied as 2x solutions to the bath at a ratio of 1:1 to avoid problems arising from slow mixing. MYC was added to the culture medium at the desired concentrations and cells were cultured for 6 and 18 h before imaging recordings. When cells were transfected with ER-GCaMP6-150, MYC was present in the culture medium containing the transfection reagents during the last 6 h before imaging recordings. Mock treatments represent the application of 0.02-0.05% v/v DMSO in bath solution or culture medium.

### Live Cell Ca^2+^ Imaging

Ca^2+^ imaging experiments were carried out in the absence of Ca^2+^ in the bath solution to prevent Ca^2+^ entry from extracellular space. Thus, the bath solution contained 140 mM NaCl, 5 mM KCl, 1 mM MgCl_2_, 0.5 mM EGTA, 10 mM glucose and 10 mM HEPES-KOH (pH 7.35). Using an iMIC microscope equipped with a polychromator V and the Live Acquisition Software (Till Photonics), we imaged cytosolic Ca^2+^ concentrations ([Ca^2+^]_cyt_) using FURA-2 (i) and Ca^2+^ concentrations in ER ([Ca^2+^]_ER_) with the Ca^2+^ sensors D1ER (ii) and ER-GCaMP6-150 (iii).

i. **Imaging cytosolic Ca**^**2+**^ **with FURA-2**. FURA-2 was excited at 340 and 380 nm alternately. The emitted fluorescence was captured at 510 nm to obtain FURA-2 images at 340 and 380 nm excitation. FURA-2 image pairs containing 40–50 cells/frame were obtained every 3 s at a magnification of 20x. FURA-2 signals were measured in FURA-2 image pairs as F340/F380, where F340 and F380 correspond to the background-subtracted fluorescence intensity at 340 and 380 nm excitation wavelengths, respectively. [Ca^2+^]_cyt_ was calculated with the standard ratiometric equation [Ca^2+^]_free_ = βK_d_,((R-R_min_)/(R_max_-R)), in which R = F340/F380 [38]. βKd, maximal and minimal F340/F380 (R_min_ and R_max_) were measured as previously described [34]. Results of FURA-2 measurements are presented as [Ca^2+^]_cyt_.
ii. **Imaging ER Ca**^**2+**^ **with D1ER**. D1ER-HEK cells were exposed to 433 nm and the emitted florescence was split at 469/23 nm and 536/27 nm to obtain the CFP and citrine components, respectively [35]. The cell fluorescence was additionally passed through a dichrotome and projected on the chip of the microscope camera to obtain simultaneous CFP and citrine images. Image pairs containing 10-15 cells/frame were obtained at 60x magnification every 10 s. The FRET ratios were calculated from background-subtracted CFP and citrine image pairs as F_Citrine_/F_CFP_, where F_Citrine_ and F_CFP_ represent the citrine and CFP fluorescence intensities at 536 nm and 469 nm, respectively [34]. Results of D1ER measurements that reflect the dynamic of [Ca^2+^]_ER_ are presented as F_Citrine_/F_CFP_ ratios.

ii) **Imaging ER Ca**^**2+**^ **with ER-GCaMP6-150**. Cells transfected with ER-GCaMP6-150 were exposed to 480 nm and the emitted fluorescence was collected at 515 nm (de Juan-Sanz et al., 2017). Images with 3-10 cells/frame were recorded at a 60x magnification every 2 s. The ER-GCaMP6-150 fluorescence (F) was measured in background-subtracted images and normalised with respect to the fluorescence measured 2 min after starting recordings (F_0_). Results of measurements with ER-GCaMP6-150 that reflect the dynamic of [Ca^2+^]_ER_ are presented as F/F_0_ ratios.

### Homology Modelling and Molecular Docking

The 476 amino acids long protein sequence of human Sec61α isoform 1 was retrieved from Uniprot (ID: P61619). Using homology modelling, structural models of human Sec61α were constructed using the following electron microscopy (EM) structures of Sec61α from *Canis lupus* (Uniport ID: P38377) as template (99.8% sequence identity using global sequence alignment): *idle* conformation (PDB code 3J7Q with resolution of 3.4 Å) [21]; *open* conformation (3JC2 with resolution of 3.6 Å) [23]; and the *inhibited* state of Sec61α in the presence of MYC (6Z3T with resolution 2.6 Å) [16]. The structural information of the missing part of 6Z3T was obtained by homology modelling based on 2WWB (EM structure with resolution 6.48 Å). Homology modelling was performed using MODELLER 9.21 [39]. After sequence alignment of target and template, MODELLER was run locally with the automodel class to generate 50 different models. As structure 3JC2 is lacking structural information for the plug region, the plug region of the open conformation was modelled as loop. The homology model with the lowest discrete optimization of potential energy (DOPE) score was selected as the final model and subjected to 1000 steps of energy minimization, using the GROMACS package (version 5.0.7) [40] to relax side chain atoms. For this, the protonation states of titratable amino acids were determined by the web server PDB2PQR [41] using the PROPKA method [42]. The total charge of Sec61α was +4e.

Two different isomers of MYC were used for docking, MYC B (E-isomer) and MYC A (Z-isomer) (see Supplementary Figure S1). Docking of MYC was conducted using AutoDock4.2 [43] to predict energetically favourable binding poses of the ligand inside or on the surface of WT human Sec61α in the idle, open and MYC-inhibited conformations. With respect to protein-ligand docking, MYC is a large compound containing 23 rotatable bonds. For each ligand, MYC A and B, the docking calculations were performed in two consecutive steps: in the first docking step, we adopted a relatively large grid box (100 Å × 100 Å × 126 Å) (see Supplementary Figure S2) covering the entire cavity of Sec61α in order to achieve an unbiased approach to determine potential binding sites. The Lamarckian genetic algorithm was employed to search for favourable binding poses, with a population size of 150, 27 × 10^3^ generations and 25 × 10^5^ energy evaluations. All other docking parameters for protein and ligand were set to the default values of AutoDock. 2000 individual docking results were clustered according to a threshold for structural similarity of 2.0 Å RMSD. In each cluster, the representative conformation was set to the one with the lowest binding free energy for that cluster. Three independent sets of 2000 docking runs each were conducted in the first stage.

In the second docking stage, the size of the grid box was reduced to 90 Å × 80 Å × 80 Å dimension (idle conformation), 80 Å × 90 Å × 80 Å dimension (intermediate conformation), and 80 Å × 80 Å × 90 Å (open conformation), respectively. This was done based on the populations of the most stable binding positions of the ligand. In the fine docking runs, more stringent parameters were used, namely 0.5 × 10^6^ generations and 100 × 10^6^ energy evaluations per run. In this stage, we executed five independent fine docking runs for the idle and intermediate conformations of Sec61α. The first step of docking to the open conformation of Sec61α showed a larger conformational diversity of MYC within the Sce61α channel pore than for the idle conformation, ten independent fine docking runs were performed for the open conformation. In each docking run we generated 50 docked conformations.

### Membrane-associated Polysome Profiling

RAW264.7 cells grown to 80% confluency on 15 cm dishes were incubated with 31.25 ng/ml MYC or DMSO carrier control for 4 h at 37°C. Membrane-associated polysomes were prepared following the methods described in Potter and Nicchitta, 2002 [44], with modifications. Cyclohexamide (CHX) (100 μg/ml) was added 5 min prior to harvesting. Cells were washed once with ice-cold PBS containing 100 μg/ml CHX then harvested in KHM buffer (25 mM K-Hepes, pH 7.2, 400 mM KOAc, 25 mM Mg(OAc)_2_, 100 μg/ml CHX). Cells were pelleted by centrifugation at 280 x g for 3 min at 4°C, then resuspended in ice-cold KHM containing 0.03% digitonin and incubated on ice for 5 min. After centrifugation again at 280 x g for 3 min at 4°C, pellets were solubilised by incubation with KHM containing 2% digitonin, 1 mM PMSF and 40 U/ml RNase inhibitor. Lysates were centrifuged at 7500 g for 10 min at 4°C and layered onto a 10-50% sucrose gradient in KMH containing 0.1% digitonin and 100 μg/ml CHX. Polysome profiling was carried out as described previously [11]. Proteins in column fractions were acetone precipitated and separated by SDS PAGE on 4-15% gradient gels (Bio-Rad) and blotted onto Immobilon PVDF membranes (Merck). Blots were probed with mouse monoclonal anti-Sec61α (Santa Cruz Biotechnology) and rabbit polyclonal anti-Sec61β antibodies [45] and detected with ECL anti-mouse IgG and anti-rabbit IgG, HRP linked antibodies (GE Healthcare), respectively.

### Statistical Analysis

Single cell data has been obtained in independent Ca^2+^ imaging recordings with 3-6 cover slips per experimental setting. Data was pooled and analysed with Excel 2010, Prism5 and Sigma Plot 10.0. The total number of analysed cells in each experimental setting is given in Figures 1-4. Statistical significance of the Ca^2+^ imaging data was assessed with the two sample Kolmogorov-Smirnov test. Statistical significance is given as n.s., non-significant; *, *p*<0.05; **, *p*<0.01 and ***, *p*<0.001.

**Figure 1.**
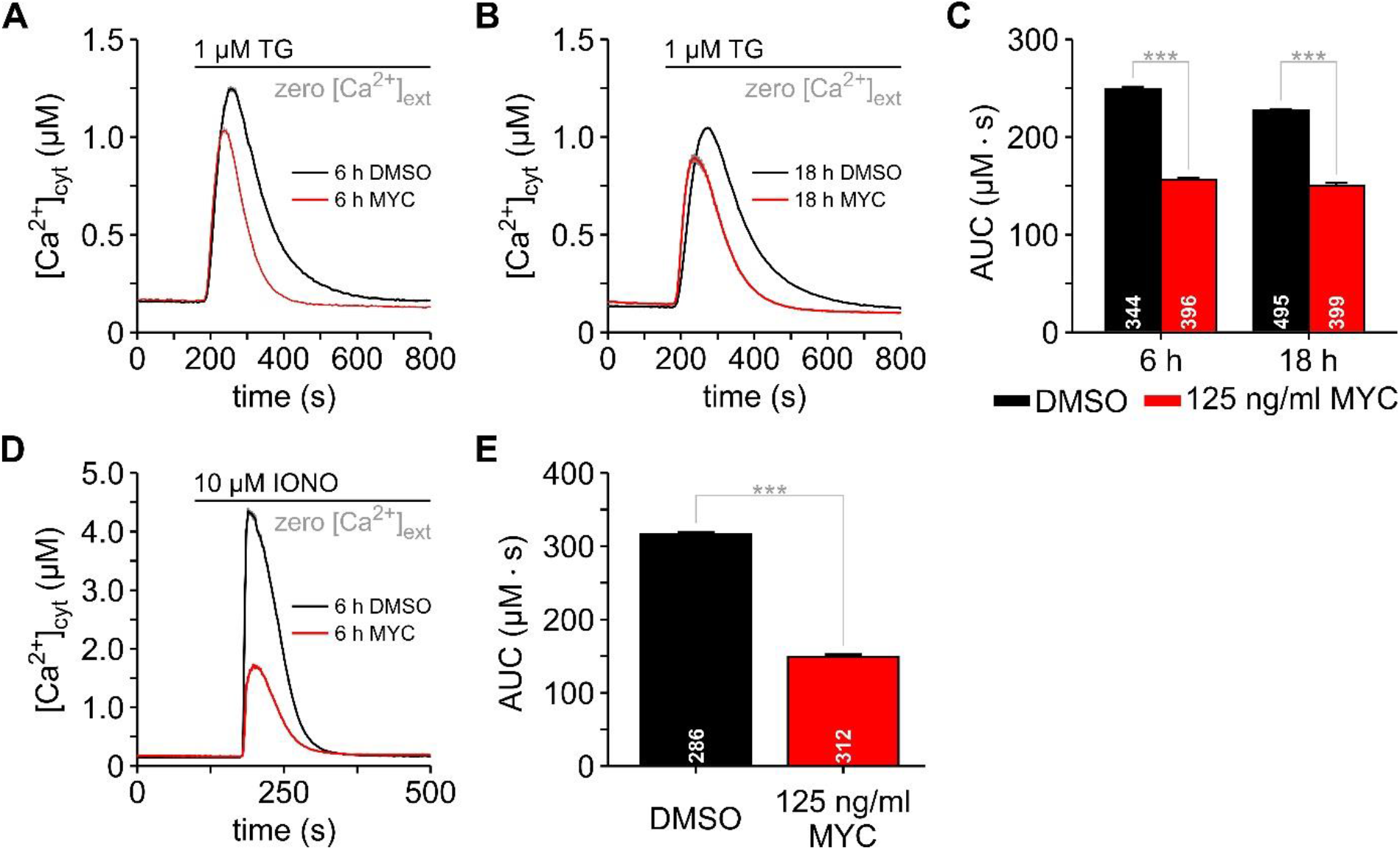
Mycolactone attenuates the Ca^2+^ mobilisation in HCT116 cells. Changes in cytosolic Ca^2+^ ([Ca^2+^]_cyt_) were imaged with FURA-2. HCT116 cells were treated with 0.05% DMSO or 125 ng/mL mycolactone (MYC) for 6 h (**A**) and 18 h (**B**) before FURA-2 loading and Ca^2+^ imaging. Thapsigargin (1 µM TG) was added to the bath solution to unmask Ca^2+^ leakage from ER (**A**-**B**). The area under the curve (AUC) of the TG-induced Ca^2+^ transients was used as a measure of the mobilisable Ca^2+^ from TG-sensitive stores (**C**). The total amount of mobilisable Ca^2+^ was estimated by treating cells with ionomycin (10 µM IONO) (**D**). The AUC of the IONO-induced Ca^2+^ transients reflect the amount of Ca^2+^ that is stored within the cells (**E**). The number of cells is given within the graph bars in **C** and **E**. Data is presented as means ± SEM; ***, *p*<0.001.

## Results

### Impact of mycolactone on the cellular Ca^2+^ homeostasis

Mycolactone has previously been shown to target the α subunit of Sec61 complexes in the ER membrane [12,16]. Biochemical evidence from translocation assays support its inhibition of Sec61α at a step after the engagement of ribosomes [18], raising the possibility that one aspect of the cellular action of mycolactone is the disruption of the Ca^2+^ homeostasis due to its interaction with Sec61α within the ER membrane. In order to explore this possibility, we analysed the Ca^2+^ homeostatic mechanisms in cells treated with mycolactone. We imaged the cytosolic Ca^2+^ concentration ([Ca^2+^]_cyt_) with FURA-2 as well as the Ca^2+^ concentration in ER ([Ca^2+^]_ER_) with the genetically encoded sensors D1ER and ER-GCaMP6-150 [34,36].

Since [Ca^2+^]_ER_ is maintained around 100-800 µM by a dynamic equilibrium between the Ca^2+^ leakage from the ER and the action of SERCA pumping Ca^2+^ back into the ER, as well as by the Ca^2+^ buffering of ER resident proteins [46,47], the SERCA-inhibitor TG unmasks the Ca^2+^ leakage producing a transient increase in [Ca^2+^]_cyt_. Typically, TG-induced Ca^2+^ transients are characterised by a rapid onset and short duration in most mammalian cells [48]. Figure 1A and B illustrates the effects of 125 ng/ml mycolactone on the Ca^2+^ mobilisation induced by TG in HCT116 cells. When Ca^2+^ transients of mycolactone-treated cells were compared with the DMSO controls, we observed a pronounced mycolactone effect on the duration and peak amplitude of the Ca^2+^ transients (Figure 1A,B). Such effects on Ca^2+^ transients are best quantified as the area under the curve (AUC), which is calculated by integration of the Ca^2+^ transients. Figure 1C shows that mycolactone significantly reduced the AUC of TG-induced Ca^2+^ transients, indicating that the amount of Ca^2+^ mobilised by TG from ER is lower in mycolactone-treated cells. However, ER Ca^2+^ depletion as well as increased Ca^2+^ leakage from ER can explain the attenuation of TG-induced Ca^2+^ transients by mycolactone. Therefore, we next used IONO in order to estimate the total amount of Ca^2+^ that can be mobilised in HCT116 cells. As shown in Figure. 1D, the IONO-induced Ca^2+^ transients had a reduced amplitude and were shorter in mycolactone-treated cells than in DMSO controls. The AUC analysis revealed that HCT116 cells lost about 2/3 of the mobilisable Ca^2+^ after 6 h treatment with 125 ng/ml mycolactone (Figure 1E). Thus, mycolactone likely compromises Ca^2+^ homeostasis in the cell by reducing Ca^2+^ storage levels, which in turn attenuates the Ca^2+^ signalling in HCT116 cells.

In HCT116 cells, less than 10 ng/ml mycolactone is sufficient to maximally reduce cell viability [15]. Since we used 125 ng/ml mycolactone in previous experiments (Figure 1), the question arises as whether low concentrations of mycolactone also induce attenuation of TG-induced Ca^2+^ transients. Therefore, we next exposed HCT116 cells to a dilution series between 3.9 ng/ml and 125 ng/ml mycolactone for 6 h. Thereafter, cells were exposed to TG to induce Ca^2+^ mobilisation from ER. As shown in Figure 2A, even 3.9 ng/ml mycolactone shortened the Ca^2+^ transients without major effect on the amplitude of the Ca^2+^ signals. Higher concentrations of mycolactone shortened the TG-induced Ca^2+^ transients further and these displayed noticeably smaller amplitudes, with the consequence that the AUC of TG-induced Ca^2+^ transients displayed a dose-dependent inhibition by mycolactone (Figure 2B).

**Figure 2.**
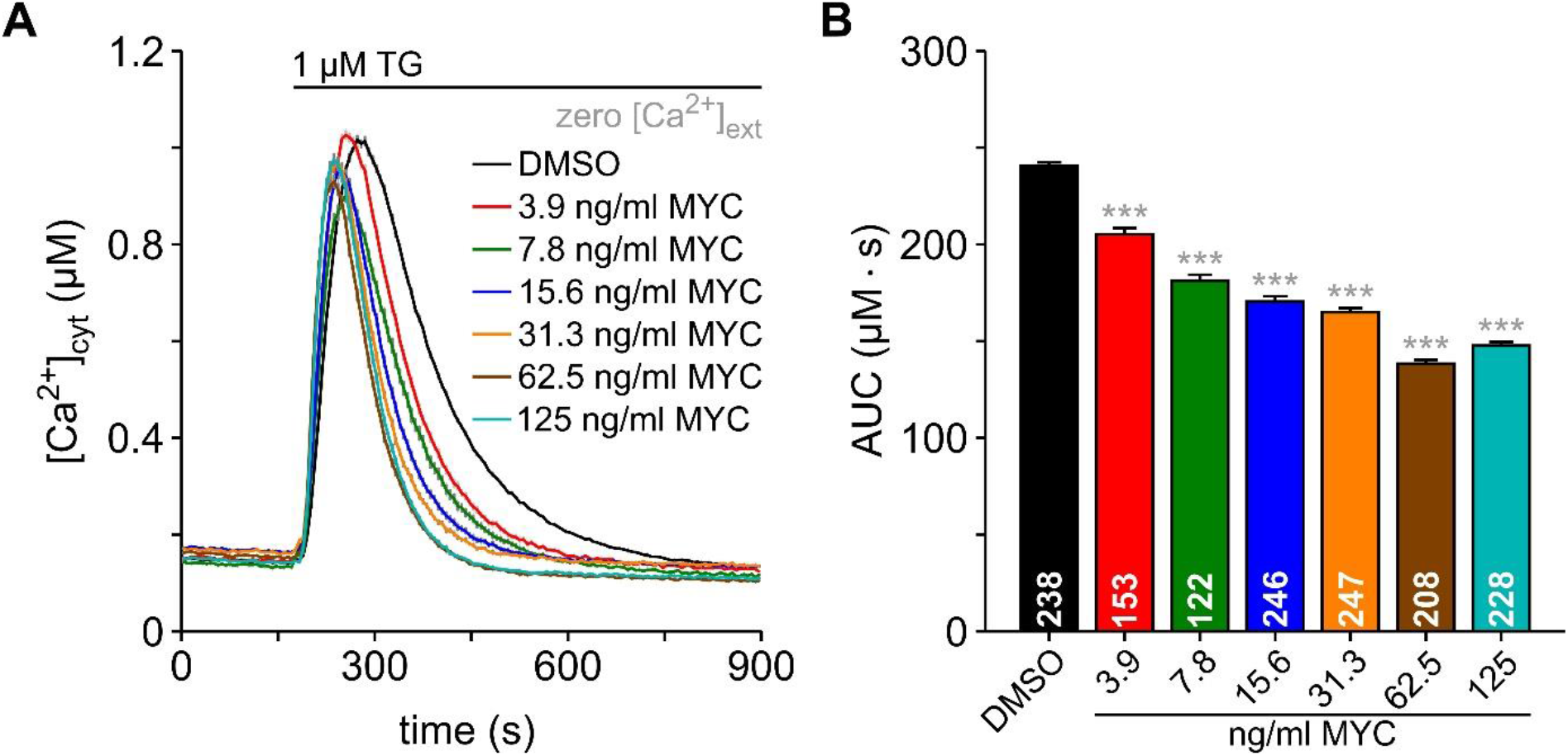
Dose-dependent effects of mycolactone on Ca^2+^ mobilisation in HCT116 cells. The dose-dependence of mycolactone (MYC) on the thapsigarin (TG)-induced Ca^2+^ mobilisation was analysed by exposing HCT116 cells to 0.05% DMSO and to a dilution series of mycolactone from 3.9 nm/ml to 125 ng/ml for 6 h. Ca^2+^ signals were imaged with FURA-2 (**A**). The area under the curve (AUC) was calculated from TG-induced Ca^2+^ transients (**B**). The number of cells is given within the graph bars in **B**. Data is presented as means ± SEM; ***, *p*<0.001.

### Mycolactone enhances the Ca^2+^ leakage from ER

The magnitude of TG-induced Ca^2+^ transients is determined by the Ca^2+^ content of the ER as well as by the intensity of Ca^2+^ leakage from ER. We found that the total amount of mobilisable Ca^2+^ in HCT116 cells is reduced by mycolactone (Figure 1D,E), suggesting that the ER Ca^2+^ content may be compromised in mycolactone-treated cells. We hypothesised that an increased Ca^2+^ leakage that leads to ER Ca^2+^ depletion during the exposure to mycolactone likely contributes to the attenuation of Ca^2+^ transients that were observed in HCT116 cells (Figure 1A-C). In order to test this hypothesis, we therefore imaged the changes in [Ca^2+^]_ER_ using the FRET-based ER Ca^2+^ sensor D1ER [35]. In experiments with ER Ca^2+^ sensors, TG typically induces an exponential depletion of ER Ca^2+^ that reflects the unmasking of the Ca^2+^ leakage from ER [34]. HEK-293 cells stably expressing D1ER (D1ER-HEK) were exposed to mycolactone for 6 and 18 h before ER Ca^2+^ imaging. As shown in Figure 3A, DMSO-treated cells display an exponential ER Ca^2+^ depletion upon application of TG. When compared with controls, the ER Ca^2+^ depletion was much faster in D1ER-HEK cells treated with mycolactone for 6 and 18 h (Figure 3A,B). Analysis of the time to 50% decay in ER Ca^2+^ (t_1/2_) revealed a two times faster decay of Ca^2+^ depletion in cells treated with mycolactone for 18 h (Figure 3C). This fast decay of Ca^2+^ depletion is indicative of a mycolactone-induced enhancement of the Ca^2+^ leakage from ER. Additionally, we observed that the basal ER Ca^2+^ levels decreased after 18 h mycolactone treatment (Figure 3B), indicating that mycolactone enhances the Ca^2+^ leakage from ER, which in the long term produces a continuous Ca^2+^ depletion in the ER.

**Figure 3.**
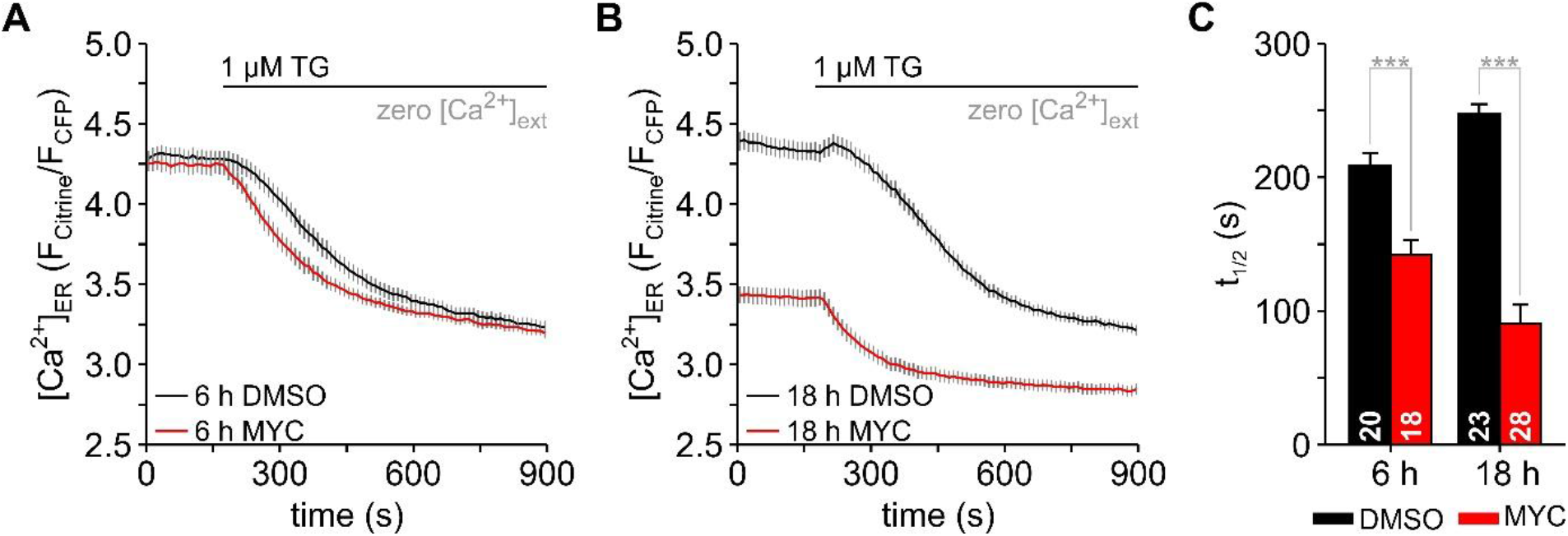
Mycolactone enhances the Ca^2+^ leakage and induces Ca^2+^ depletion in the ER. Changes of the Ca^2+^ concentration in ER ([Ca^2+^]_ER_) were imaged with the FRET-based D1ER sensor and are expressed as F_Citrine_/F_CFP_ ratios. HEK-293 cells expressing D1ER (D1ER-HEK) were exposed to mycolactone (MYC) for 6 and 18 h (**A**, 6 h mycolactone, 100 ng/ml. **B**, 18 h mycolactone, 125 ng/ml). DMSO was used as a control (**A**, 0.020 %. **B**, 0.025 %). Thapsigargin (1 µM TG) was applied to unmask the Ca^2+^ leakage form ER. The mycolactone exposure of 6 h produced a faster decrease of [Ca^2+^]_ER_ after TG application, indicating an enhancement of Ca^2+^ leakage from ER (**A**). In cells that were exposed to mycolactone for 18 h, an additional decrease of basal [Ca^2+^]_ER_ was observed, which is indicative of Ca^2+^ depletion in ER (**B**). The speed of TG-induced changes in [Ca^2+^]_ER_ was measured as the time that D1ER ratios needed to decrease to 50 % (**C**, t_1/2_). The number of cells is given within the graph bars in **C**. Data is presented as means ± SEM; ***, *p*<0.001.

### Sec61 complexes are involved in the mycolactone effects on ER Ca^2+^ leakage

Since mycolactone targets the α subunit of Sec61 complexes in the ER membrane [12,16] and Sec61 complexes form a Ca^2+^ leakage pathway from ER [49], a distinct possibility is that mycolactone stabilises a conformation, in which Sec61 complexes are open and permeable for Ca^2+^. To test this hypothesis, we took advantage of a series of mycolactone-resistant HCT116 cell clones that have heterozygous non-synonymous mutations within the *SEC61A1* gene encoding Sec61α [15,16]. For our Ca^2+^ imaging experiments, we selected the cell line expressing Sec61α D60G, which is located in the cytosolic loop (CL) 1 of Sec61α, as well as the cell lines expressing Sec61α S82Y and Q127K, which are located in the transmembrane segments (TM) 2 and 3, respectively, and the mutations R66K and S71F which are located in the transition between CL1 and TM2.

Using the ER Ca^2+^ sensor ER-GCaMP6-150 [36], we have visualised the TG-induced depletion of ER Ca^2+^ as a surrogate of Ca^2+^ leakage in these cell lines. As expected, ER-GCaMP6-150 detected an exponential Ca^2+^ depletion in wild type (WT) HCT116 cells exposed to DMSO for 6 h (Figure 4A), as well as in all clones containing Sec61α mutants (Figure 4B-F). Notably, the rate of Ca^2+^ depletion was different in each clone as revealed by the t_1/2_ values (Figure 4G), particularly for D60G and S82Y. This suggests that the mutations themselves may impact the structure and function of Sec61 translocons, as expected as they are *prl* mutants.

**Figure 4.**
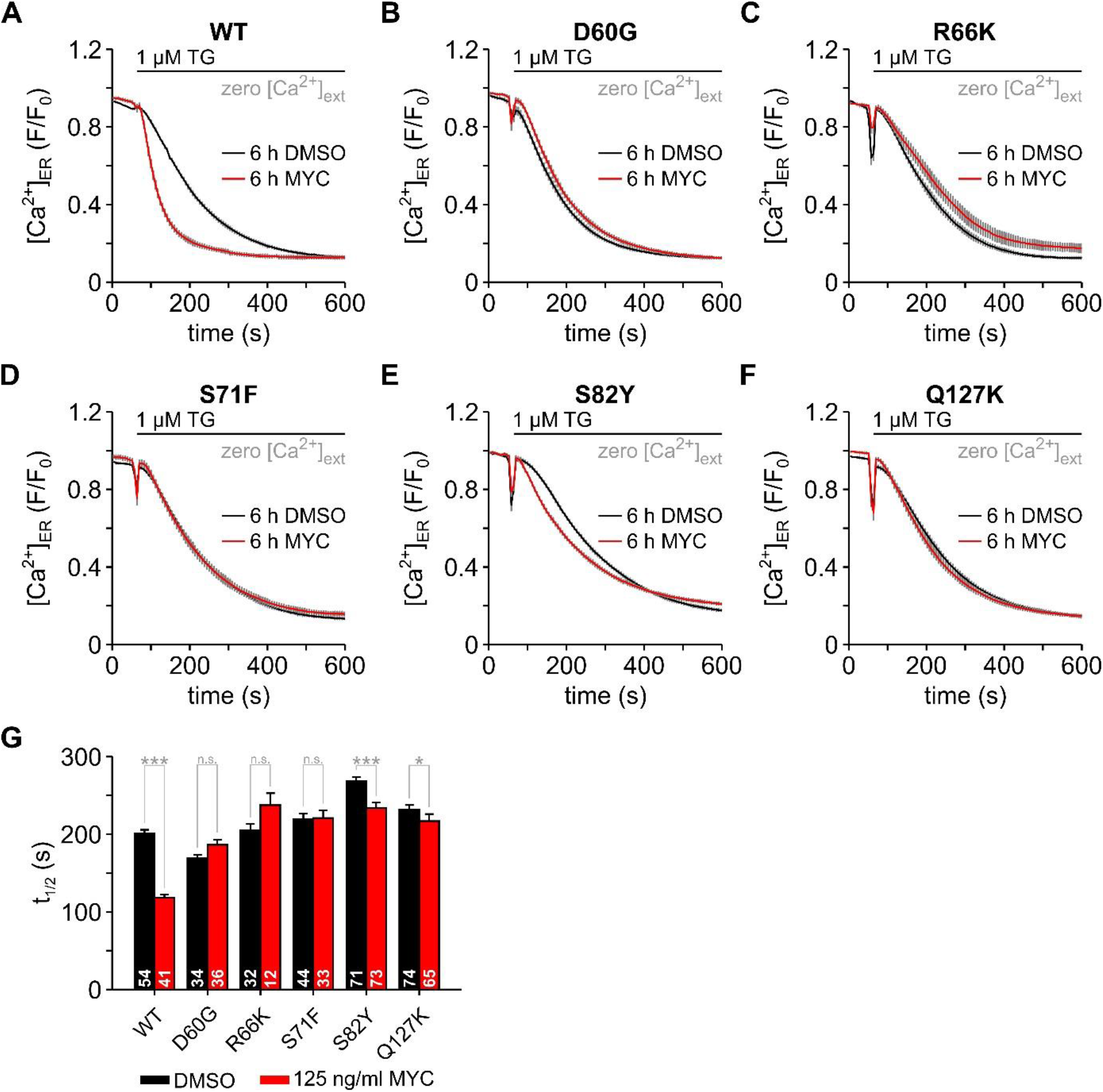
Mycolactone-resistant mutations in Sec61α abolish the effects of mycolactone on the Ca^2+^ leakage from ER. The effects of mycolactone (MYC) on [Ca^2+^]_ER_ were analysed in wild type HCT116 cells (**A**, WT) and in HCT116 cells containing point mutations in Sec61α (**B**, D60G; **C**, R66K; **D**, S71F; **E**, S82Y; **F**, Q127K). In order to image [Ca^2+^]_ER_, cells were transfected with the ER Ca^2+^ sensor ER-GCaMP6-150. Changes in [Ca^2+^]_ER_ are given as normalised fluorescence (F/F_0_). Cells were treated with 0.05% DMSO or 125 ng/ml mycolactone for 6 h before Ca^2+^ imaging. Thapsigargin (1 µM TG) was used to induce ER Ca^2+^ depletion. Accordingly, the speed in the decrease of [Ca^2+^]_ER_ is a measure of the Ca^2+^ leakage from ER and was calculated as the time to 50% decrease in the normalised fluorescence F/F_0_ (**G**, t_1/2_). The number of cells is given within the graph bars in **E**. Data is presented as means ± SEM; *, *p*<0.05; ***, *p*<0.001.

In WT HCT116 cells mycolactone reduced the t_1/2_ by about 50% (Figure 4A,G), in line with the observation using the ER Ca^2+^ sensor D1ER (Figure 3C). However, the effects of mycolactone on cells expressing the Sec61α mutants varied slightly according to the mutation. For instance, mycolactone no longer caused any change to the t_1/2_ with the Sec61α S71F mutant (Figure 4D,G). Although not statistically significant, longer t_1/2_ values were observed in the Sec61α D60G and Sec61α R66K mutants (Figure 4B,C,G). Compared with the respective DMSO controls, significantly shorter t_1/2_ values were detected in the Sec61α mutants S82Y and Q127K (Figure 4E-G), although these mycolactone effects were always less pronounced than the nearly 50 % reduction of t_1/2_ in WT HCT116 cells. In the case of the Sec61α S82Y mutant, mycolactone reduced t_1/2_ by about 12% (Figure 4E,G), a much milder effect. All in all, the direct analysis of the TG-induced ER Ca^2+^ depletion provided evidence that mycolactone no longer enhances Ca^2+^ leakage from ER in HCT116 cells expressing the mycolactone resistance mutations in Sec61α (Figure 4). Hence, our results strongly support the suggestion that Ca^2+^ leakage from ER in the presence of mycolactone is mediated by Sec61α. Supporting this suggestion, siRNA knockdown of *SEC61A1* in HeLa cells abolished the enhancing effects of mycolactone on Sec61α-mediated Ca^2+^ leakage (data not shown). Furthermore, the absent or weakened Ca^2+^ leakage from the ER could be a contributing factor to the increased mycolactone resistance of these cell lines in cell survival assays [15,16].

### Molecular Docking

For our homology model of the *idle* conformation of human Sec61α within ribosome-Sec61 complexes, docking yielded favourable low-energy conformations where mycolactone A and B interacted with Sec61α with a binding energy of −7.25 ± 0.8 kcal/mol and −8.17 ± 0.5 kcal/mol (on average), respectively. Both mycolactone isomers were placed near the cytosolic entrance of the *idle* WT Sec61α pore (see Figures 5 and 6, bottom left panels) and mainly occupied the volume between cytosolic loops 6 and 4 (CL6 and CL4). The long acyl tails (Southern tail) of both mycolactones face towards the CL4 region. The docking results suggest that the Northern acyl side chain of both mycolactone A and B could potentially form hydrogen bonds with the CL6 region because residues from the region Pro280-Ser287 (CL6-TM7(transmembrane helix 7)) were predicted to strongly interact with both isomers of mycolactone.

**Figure 5.**
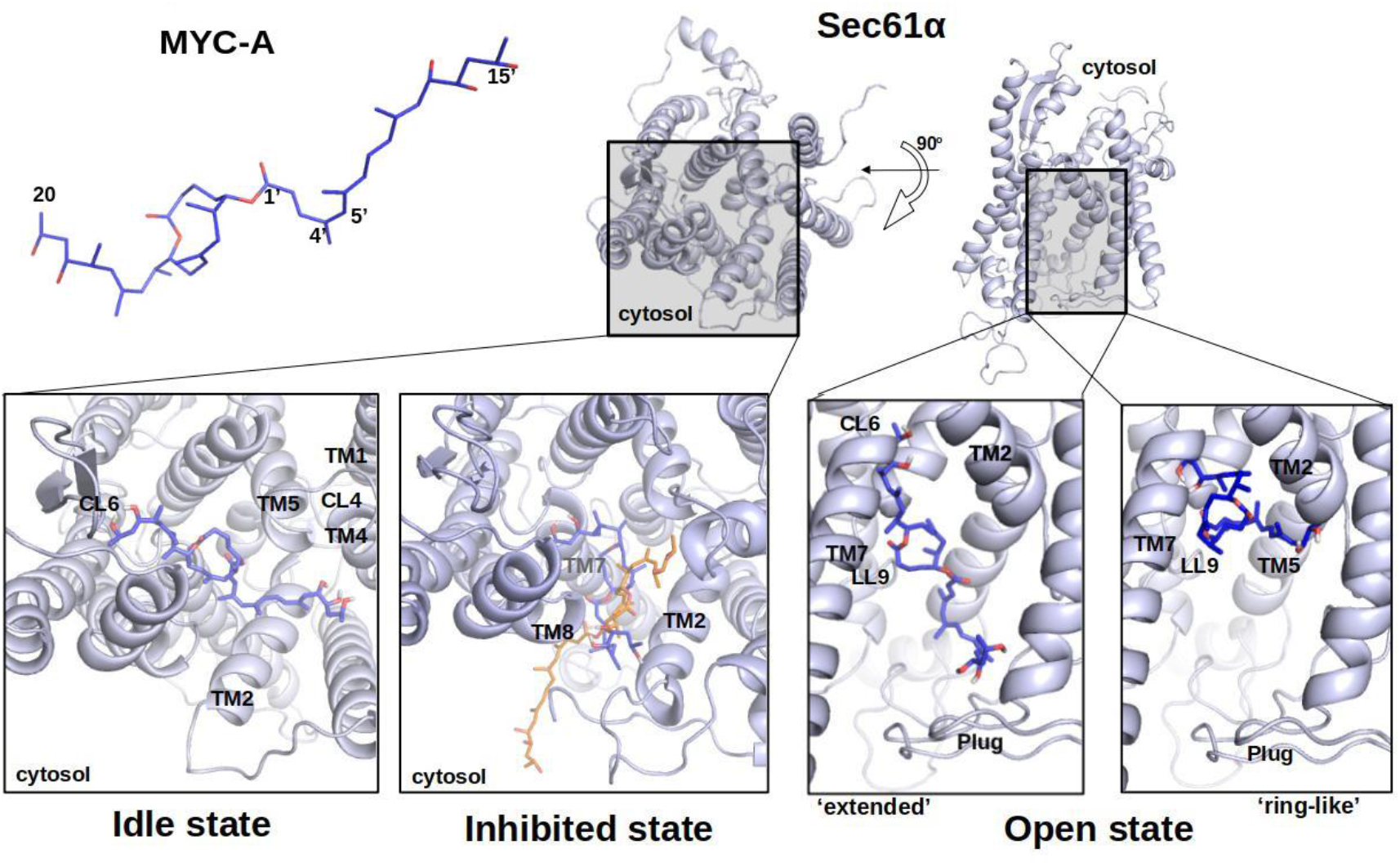
Most favourable binding poses of mycolactone A (MYC-A, blue) in *idle* (lower-left), *mycolactone-inhibited* (lower-middle) and *open* (lower-right) state-models of human WT-Sec61α. The mycolactone conformation from cryo-EM structure 6Z3T is super imposed on the docking result (orange, “inhibited state”).

**Figure 6.**
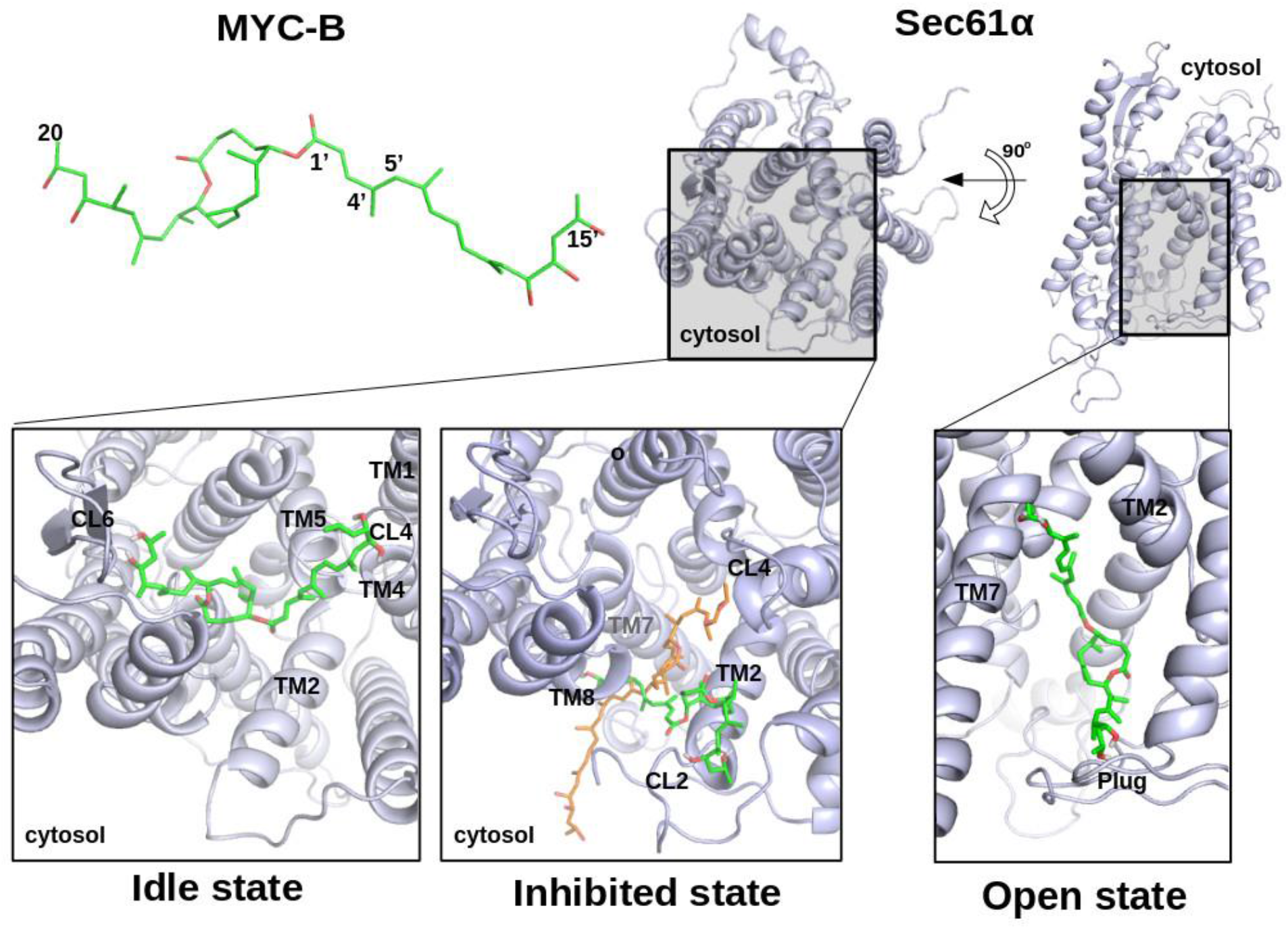
Most favourable binding poses of mycolactone B (MYC-B, green) in *idle* (lower-left), *inhibited* (lower-middle) and *open* (lower-right) state-models of human WT-Sec61α. The mycolactone conformation from cryo-EM structure 6Z3T is super imposed on the docking result (orange, “inhibited state”).

For the mycolactone-inhibited conformation, the docking results suggest that mycolactone occupies the groove between TM8, TM2 and TM7 helices (Figures 5 and 6, bottom middle panels). The docking position of the lactone core of mycolactone compares well to the orientation of mycolactone determined in the EM structure [16]. In contrast to the EM structure, the longer acyl side chain (southern chain) of mycolactone is oriented towards the lateral gate (Figures 5 and 6, bottom middle panels), possibly due to the absence of lipids in the docking analysis. These docking results suggest that both mycolactones A and B can interact with this conformation of Sec61α with similar binding energies of −8.21 ± 1.1 and −8.67 ± 0.6 kcal/mol, respectively. Residues from the region Leu89-Lys98 (TM2) strongly interact with both isomers of mycolactone. The most favourable final binding poses of mycolactone A and B are displayed in Figures 5 and 6.

For the *open* conformation of Sec61α, the docking results placed mycolactone inside the channel pore that would normally be the translocation path of substrate peptides. Mycolactone A was placed here in two main arrangements that we refer to as ‘ring-like’ or ‘extended’ conformations (Figure 5, bottom right panel). The ‘extended’ conformation was slightly more favourable with a binding energy of −9.02 ± 0.5 kcal/mol than the ‘ring-like’ conformation (−8.21 ± 0.6 kcal/mol). In the ‘extended’ conformation, the acyl side chains of mycolactone A often formed hydrogen bonds with residues in CL6, lumenal loop (LL) 9 and the plug region. In the ‘ring-like’ conformation, mycolactone A interacted with the N-terminal region of TM2, LL9 and TM5. mycolactone B also adopted the ‘extended’ conformation (see Fig. 6) inside the channel pore and the ligand pose looked like a ‘bridge’ between the plug region and the loop CL6 similar to mycolactone A. The acyl side chains of mycolactone B often interacted with residues of CL6, LL9, plug and TM5. The lowest binding energy of mycolactone B was −8.78 ± 0.6 kcal/mol.

### Functional consequences of Sec61α blockade by mycolactone

Previous studies have shown that mycolactone causes an eIF2α-dependent reduction in overall protein translation rates, which reduces the abundance of polysomes and increases that of the 60S ribosomal subunit peak in the sub-polysomes [11], albeit more mildly and more slowly than strong inducers of the integrated stress response, such as tunicamycin. Since puromycin, which disassembles ribosomal subunits leading to loss of polysomes [50], increases the Sec61-mediated Ca^2+^ leakage from ER [51], we next analysed the effects of mycolactone on polysomes. Polysome profiling is typically carried out in 1% triton cell extracts, where cytosolic ribosomes predominate. In order to investigate the effect of mycolactone on polysomes at the ER, we analysed the membrane-associated polysome profile of cells exposed to mycolactone using digitonin extracts. This showed that mycolactone reduced the polysomal fractions and increased the 80S peak (Figure 7A). In addition, immunoblotting of the polysome gradient fractions showed a shift of Sec61α from the polysomal fraction in control cells to the 80S fraction in mycolactone-treated cells (Figure 7B). This data suggests that the addition of mycolactone results in the disassembly of polyribosomes leaving isolated ribosome-Sec61 complexes in the ER membrane. Furthermore, it confirms previous data from extracted canine microsomes that mycolactone appears not to displace ribosomes from the ER [16,18].

**Figure 7.**
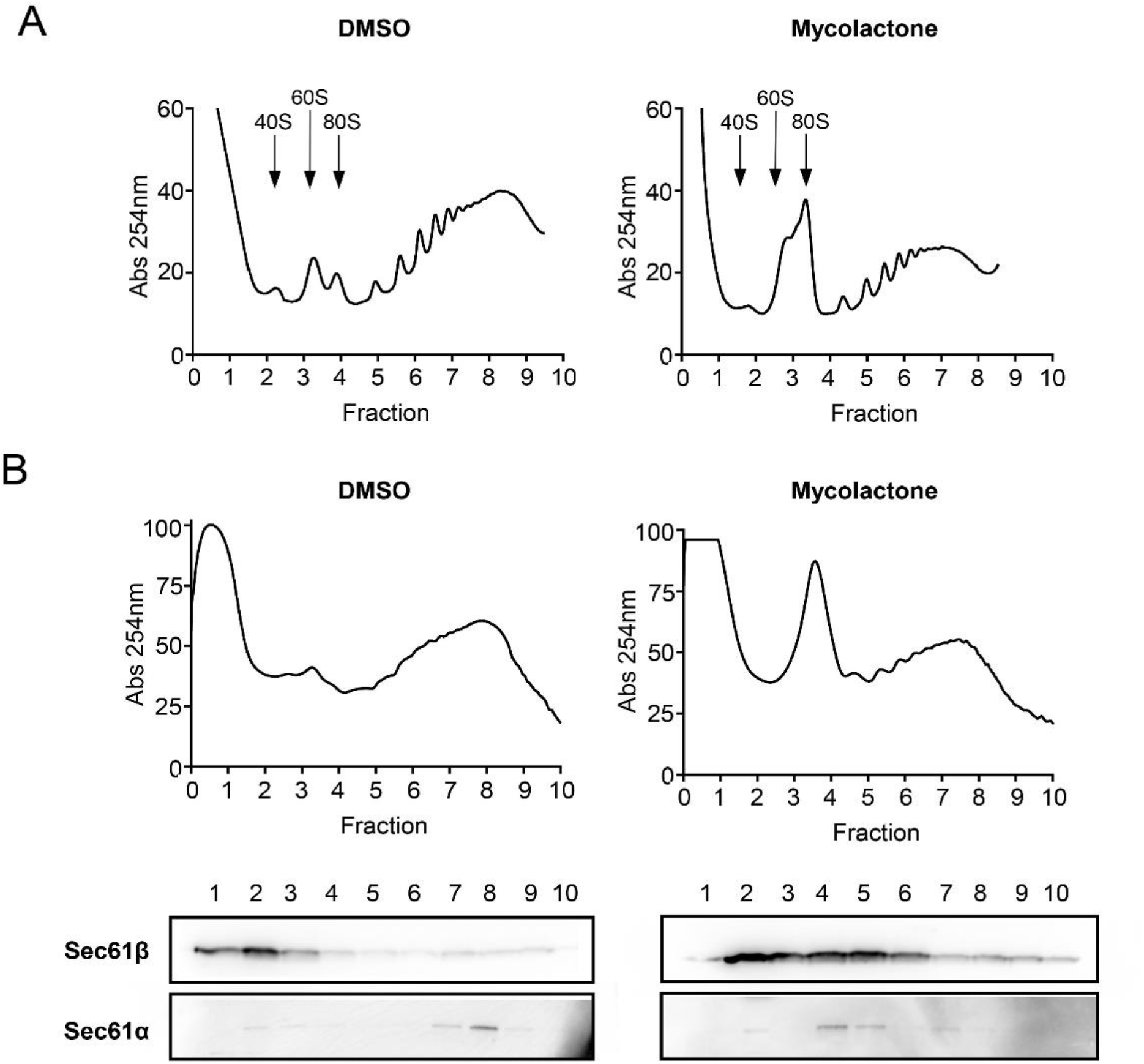
Mycolactone promotes the disruption of polysomes. Typical polysome profile of whole cell triton-x-100 extracts from RAW264.7 cells treated with DMSO or 31.25 ng/ml mycolactone (**A**) and of membrane associated polysomes from RAW264.7 cells treated with DMSO or 31.25 ng/ml mycolactone for 4 h (**B**). Membrane associated polysomes were isolated from semi-permeabilised cell extracts solubilised with 2% digitonin and separated on a 10-60% sucrose gradient as described in Material and Methods. Western blotting of column fractions shows translocon components associated with the 80S peak in digitonin solubilised RAW264.7 cell membrane fractions after mycolactone treatment (**C**). Blots were probed with anti-Sec61α and anti-Sec61β antibody. This figure is representative of duplicate assays.

## Discussion

In this work, we present novel data from Ca^2+^ imaging and show how application of the Sec61α inhibitor mycolactone affects Ca^2+^ leakage through Sec61α. Also, we present results from molecular docking aimed at identifying putative binding modes of mycolactone to various conformational states of Sec61α. We will start this discussion section by summarizing the existing experimental data on how mycolactone affects Sec61α function. Then, we will connect this with the available structural data and docking results and, finally, suggest an integrated mechanistic model for mycolactone action on Sec61α.

It has been known for some time that Sec61α can facilitate the Ca^2+^ leakage from the ER to the cytosol [51], which is normally pumped back into the ER by the SERCA pump. Application of the SERCA pump inhibitor TG prevents this and leads to a complete emptying of the ER Ca^2+^ due to the Ca^2+^ leakage. This is followed by partitioning of Ca^2+^ to other compartments or release from cells. The time course of the TG-induced spike in [Ca^2+^]_cyt_ is represented by the black curves in Figure 1. Application of mycolactone decreases the peak and duration of this Ca^2+^ release signal (red curves in Figure 1), hence indicating that the ER membrane becomes more permeable for Ca^2+^ when mycolactone binds. The titration experiments shown in Figure 2 demonstrate that these effects on Ca^2+^ leakage increase dose-dependently with increasing mycolactone concentration, presumably as the available Sec61 translocons become occupied with mycolactone, since their stoichiometry is approximated 1:1 [16]. Monitoring the [Ca^2+^]_ER_ (Figure 3) revealed that pre-incubation with mycolactone led to a faster Ca^2+^ release from the ER than in the control, as expected based on [Ca^2+^]_cyt_ measurements. Extending the pre-incubation time with mycolactone from 6 to 18 h led to a strong Ca^2+^ depletion in ER, presumably due to increased leakage that SERCA could no longer control. Further analyses of the Ca^2+^ depletion in HCT cells carrying mycolactone-resistant Sec61α mutations enabled us to pin down the role of Sec61 translocons in the effects of mycolactone on the Ca^2+^ leakage from ER (Figure 4). We observed no statistically significant effects of mycolactone with the Sec61α mutants D60G, R66K and S71F, indicating that mycolactone has no effect on the Ca^2+^ leakage supported by these Sec61α mutants.

Mycolactone induced a slight increase of Ca^2+^ leakage only in cells expressing the mutants S82Y and Q127K, but these effects were by far less pronounced than that in control cells. This demonstrates that the effect of mycolactone on Ca^2+^ leakage is extremely likely to be related to its interaction with Sec61α rather than its other reported targets, or a general disturbance of phospholipid bilayer integrity. The effects of mycolatone on the Ca^2+^ leakage mediated by the S82Y and Q127K mutants may reflect differences in the effect of the mutations on the structure and function of the Sec61 translocon. This is not unexpected, since most have been previously associated with the *prl* phenotype, which alters the efficiency of translocation depending on the nature of the signal peptide. Further Ca^2+^ imaging experiments are required to characterise the kinetic of the Ca^2+^ leakage mediated by mycolactone-resistant Sec61 mutants. McKenna et al. (2016) found that ribosomes still bind to the Sec61 complex in the presence of mycolactone but observed discrete changes in the architecture of the ribosome– nascent chain (RNC)-Sec61 interaction, as evidenced by cross-linking of the nascent chain to Sec61 subunits [18]. Additionally, mycolactone treatment altered the trypsin sensitivity of CL6 and CL8 (implicated in ribosome binding) of Sec61α. On this basis, this biochemical analysis favoured a model where mycolactone perturbs the interaction between signal peptide and the ribosome-Sec61 complex that is necessary for co-translational translocation to progress. The recent analysis of the mycolactone-bound translocon shows that the conformation of the translocon is not long similar to that of the *idle* translocon, with Sec61α instead adopting a position that is wedged open at the lateral gate [16]. The lactone ring of mycolactone binds near the lateral gate and stabilises it in a partially open conformation.

Molecular re-docking of mycolactone into a homology model of the EM structure of the mycolactone-bound state confirms the plausible position of the lactone ring identified by Gérard et al. (2020). In these docking runs, the hydrophilic chains bound inside the Sec61α pore, presumably because the lipid bilayer was not present as an alternative hydrophobic surface. Previously, Aydin et al. (2019) characterized the orientational preference of mycolactone in a plain DPPC bilayer by molecular dynamics simulations [52]. They reported that the highly hydrophobic lactone ring prefers to be buried among the hydrophobic lipid tails of the phospholipid bilayer, whereas the hydroxyl groups that are found on the side chains either tend to stretch out to the hydrophilic lipid headgroups or interact with water molecules. Hence, the more extended conformation of mycolactone reported by Gérard et al. (2020) appears more plausible than the compact conformation identified in the docking run [16]. Molecular docking of mycolactone to the *idle* state also revealed a novel potential binding site on the surface of Sec61α (discussed in detail below). Molecular docking of mycolactone to the open state identified further putative binding positions in the channel pore. However, noting the just mentioned preference of mycolactone’s lactone ring and its Northern and Southern chains for a patterned, mixed hydrophobic/hydrophilic environment [52], these docking positions in the water-filled hydrophilic pore of the open state appear less likely. Also, there is so far no experimental evidence supporting such binding modes.

Similar to Gerard et al. (2020), these docking runs position mycolactone with the core and Northern chain within the translocon and the Southern chain extending from the translocon into the ER membrane. It has been known for some time that the Southern chain of mycolactone is important for cytotoxic and/or cytokine suppressing activity of mycolactone [3]. Since forward genetic screens have unambiguously linked these readouts to Sec61α [12,15,16], this data can be used as a retrospective surrogate for Sec61α inhibition. Here, naturally occurring mycolactones with differing specific activities are invariant in the Northern chain and lactone core, but vary in the Southern chain methylation, hydroxylation and/or length. Furthermore, wide-ranging structure-function studies using synthetic variants of mycolactone [53] have revealed that variations in the Northern chain are generally well-tolerated, whereas the Southern chain is much more sensitive to even small perturbations in structure. After addition of even bulky functional groups to the Northern chain at C14 or C20 mycolactone retains cytotoxic activity. On the other hand, truncation of the Southern chain to C6′ or C2′, or completely removing it results in a molecule with much reduced activity. A synthetic mycolactone produced by Scherr et al. (2013), with no hydroxylation of the Southern chain, was found to retain minimal cytotoxic activity [54]. More specifically to translocation, structural variants of mycolactone lacking the Northern chain can still compete with cotransin (CT7) for Sec61α binding, whereas those lacking the Southern chain or both chains cannot [12]. These data strongly suggest that the lactone core and Northern chain interacts directly with TM2 of the translocon, explaining the evolutionary conservation of these elements of the structure. On the other hand, we propose that the Southern chain has essential interactions with the lipid bilayer and ascribe the activity differences in Southern chain variants to differing abilities to interact with the lipid bilayer. Interactions of the tail groups with phospholipid headgroups may serve to anchor the molecule, stabilising the inhibited conformation of the translocon.

The *idle* state represents a non-translocating ‘primed’ state of Sec61α bound to ribosomes with closed lateral gate and the plug in place. This state is represented by the structure reported by Voorhees et al. (2014) and, so far, there is no evidence that Sec61 translocons support Ca^2+^ leakage in this *idle* state [55]. After binding of the ribosome and shortly before engagement of the signal peptide within the lateral gate, we postulate that Sec61α rapidly transitions between this and an *intermediate* state, that closely resembles that of the yeast Sec62/Sec63-bound translocon required for post-translational translocation in that species. Bhadra et al. (2021) recently characterized by molecular dynamics simulations how Sec63 modulates opening of the lateral gate of Sec61 in yeast [56]. However, this transient structure has not yet been seen in the absence of mycolactone or other inhibitors [16]. It is plausible that the conformation of the *intermediate* state in Figure 8 may differ from that reported for the mycolactone-Sec61α complex, which has an open lateral gate, and the plug helix is intact but displaced. It is conceivable that “translocon breathing” could be associated with very transient states in which Ca^2+^ ions can leak out of the ER. If so, mycolactone’s stabilisation of this state would clearly support the leak of Ca^2+^ ions observed even at doses where all translocons are expected to be saturated with mycolactone.

**Figure 8.**
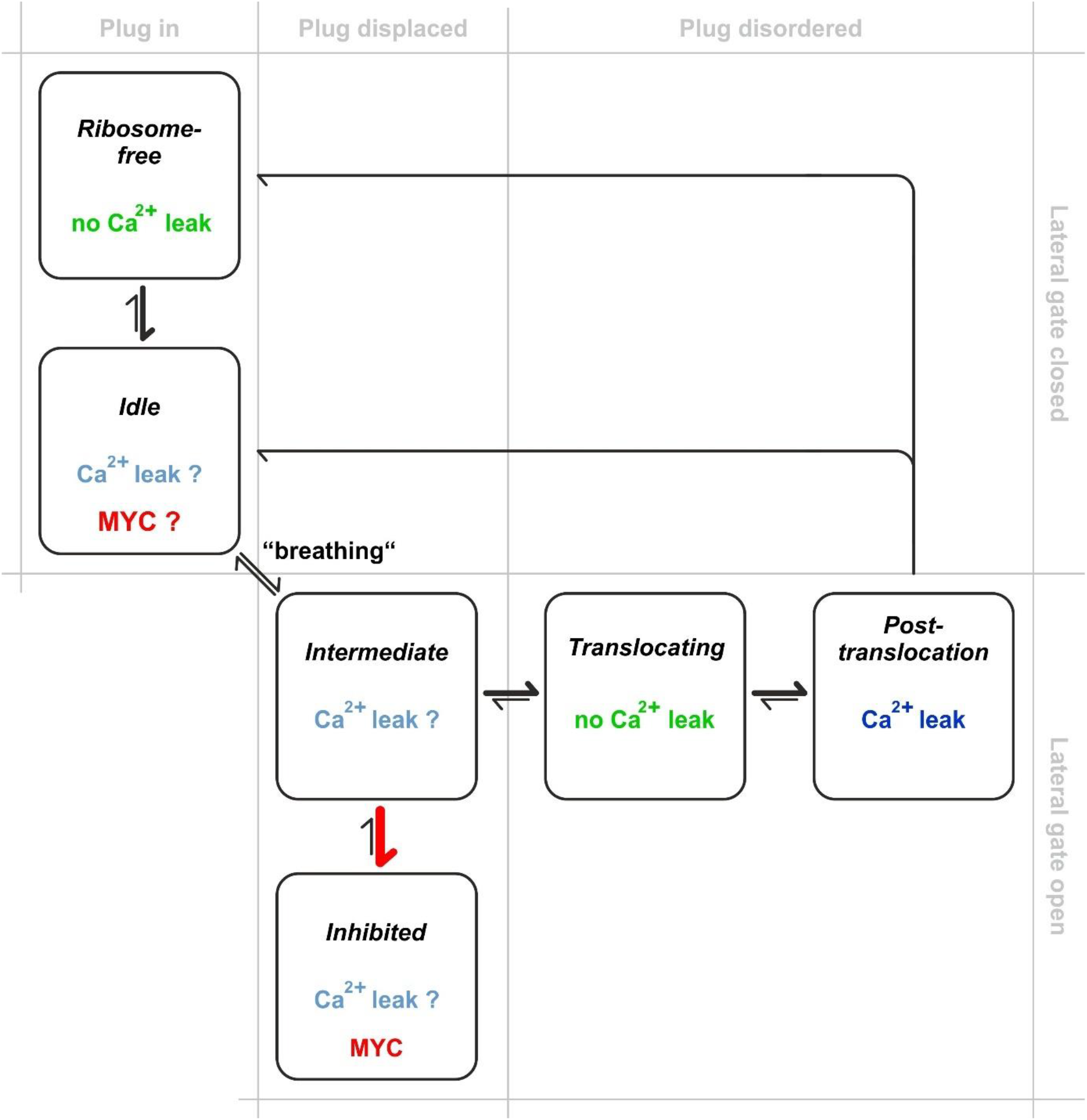
Model for mycolactone (MYC)-induced Ca^2+^ leak via Sec61 complexes. In normal translocation, *idle* translocons are engaged with ribosome nascent chain complexes, but co-translational translocation has not yet started. There is no evidence that *idle* translocons are Ca^2+^ permeable. Translocons “breathe” along the lateral gate, rapidly transitioning between the *idle* and *intermediate* states. We propose that the *intermediate* state may become transiently Ca^2+^ permeable due to its open structure and the partial displacement of the plug helix. Once the signal peptide of the nascent chain functionally engages with the lateral gate of Sec61α, the protein-conducting channel facilitates the translocation of the nascent protein by complete unfolding of the plug into a disordered state, but the *translocating* state is not permeable to Ca^2+^ due to the peptide filling the resultant channel. After translocation, Sec61 complexes transiently remain in this fully open *post-translocation* state, but without a nascent chain, allowing Ca^2+^ ions to permeate through the aqueous channel. The cycle is completed when the ribosomes de-attach and Sec61 complexes return to the *ribosome-fre*e or *idle* state, depending on whether the ribosome is released. Mycolactone traps translocons in an *inhibited* state, that resembles the *intermediate* state, but may also be able to bind Sec61α in the *idle* state. This locks the Sec61 translocon in a Ca^2+^-permeable conformation and an inefficient cycling through the other states, resulting in enhanced Ca^2+^ leakage from the ER. Mycolactone causes an increase in membrane-associated 80S ribosomal monosomes. Hence, another contributing factor to the enhanced Ca^2+^ leakage could be sustained dwell time in the *post-translocation* state, due to decreased efficiency of ribosome release.

Considering the known three conformations of Sec61α (*idle, intermediate, open*), we postulate that mycolactone preferentially binds to, and stabilises, the *intermediate* conformation, which is structurally similar to the *inhibited* conformation. Translocon breathing suggests a mechanism for mycolactone’s accessing of the translocon, as unbound mycolactone distributed in the ER membrane could penetrate the lateral gate during this step. This is in line with the structure–activity relationship data for mycolactone variants [54,57,58]. Alternatively, our docking analysis predicts that mycolactone could potentially bind on the cytosolic entrance of the Sec61α pore.

In the scheme of Figure 8, nascent chain translocation occurs after the plug has moved out of the pore (*translocating* state). Since the pore is entirely occupied by the translocating peptide, it is not thought that there is Ca^2+^ leakage in this state. When translocation is completed, the channel pore is briefly left open (*post-translocation* state) by the exiting peptide making this state Ca^2+^ permeable. Finally, the lateral gate and plug close again and the channel converts back to the *idle* state or *ribosome-free* state depending on whether the ribosome detaches from the translocon or not. While there is no evidence that Ca^2+^ ions permeate through Sec61 translocons in the *idle* state[55], functional studies in planar lipid bilayers demonstrated that Sec61 translocons support Ca^2+^ leakage in the *post-translocation* state but not in the *translocating* and *ribosome-free* states[51,55,59].

The breakdown of the polysomes to a single 80S peak hints at effects of mycolactone on this stage of translocation that may also contribute to the Ca^2+^ leakage. While translocation of secreted and Sec61α-dependent single-pass membrane proteins is blocked by mycolactone [60], many multi-pass membrane proteins are still being synthesised, not all of which are thought to be substrates of the ER membrane protein complex [61]. We found that monomeric ribosomes accumulate in cells treated with mycolactone, and these are still associated with Sec61α. This scenario could be explained by altered conformation of the RNC complex delaying ribosome release following translation termination. Alternatively, the loss of the polysomes may slow the transition between the *post-translocation* and *idle/ribosome-free* states, allowing more time for Ca^2+^ to leak out. Interestingly, this data is in line with a previously published study, which showed a similar effect in cells exposed to DTT or TG [62]. Like mycolactone, these compounds activate the integrated stress response pathways, and here the monosomal ribosomes continued to translate while associated with the membrane. Moreover, it agrees with our docking analysis that suggests that mycolactone can form hydrogen bonds with residues of CL6 on Sec61α, which interact directly with the ribosome. We speculate that the interaction of mycolactone with CL6 could change the conformation of these regions and consequently could alter the interaction and/or rate of release. This fits well with the changes in trypsin sensitivity of CL6 and CL7 observed by McKenna et al. (2016). Other factors in the cellular response to mycolactone may further impact Ca^2+^ homeostasis. For example, the gradual loss of ER chaperones, such as BiP, as well as oxidoreductases and other Ca^2+^ regulators could also lead to enhanced leakage [14].

In summary, binding of mycolactone to Sec61α is reminiscent of the “foot-in-the-door” mechanism suggested earlier for eeyarestatin compounds [34]. By stabilising the translocon in an *intermediate* state that is barely capable of Ca^2+^ leakage and/or altering the efficiency with which the translocon can cycle through its various structural states, mycolactone promotes an enhanced Ca^2+^ leakage from the ER that likely underpins its cytotoxic effects.

## Data availability

All supporting data in relation to the studies reported here are provided in this manuscript.

## CRediT Contribution

**Pratiti Bhadra:** Investigation, Data curation, Formal analysis, Writing – review and editing. **Scott Dos Santos:** Investigation, Data curation, Formal analysis. **Igor Gamayun:** Investigation, Data curation, Formal analysis. **Tillman Pick:** Formal analysis, Writing – review and editing. **Joy Ogbechi:** Investigation, Formal analysis. **Belinda S. Hall:** Investigation, Formal analysis, Writing – review and editing. **Richard Zimmermann:** Conceptualization, Supervisor, Funding acquisition, Writing – review and editing. **Volkhard Helms:** Conceptualization, Supervisor, Funding acquisition, Writing – original draft, Writing – review and editing. **Rachel E. Simmonds:** Project administration, Conceptualization, Funding acquisition, Writing – review and editing. **Adolfo Cavalié:** Project administration, Conceptualization, Funding acquisition, Writing – original draft, Writing – review and editing.

## Acknowledgements

We would like to thank Dr Yoshito Kishi (Harvard University, USA) for the gift of synthetic mycolactone A/B. We thank Roger Y. Tsien for kindly providing us the calcium sensor D1ER and to Jaime de Juan-Sanz and Timothy (ICM-Institut du Cerveau, Paris, France) and A. Ryan (Weill Cornell Medicine, New York, USA) for the ER-GCaMP6-150 construct, respectively. We acknowledge Heidi Löhr and Martin Simon-Thomas (Saarland University) for their excellent technical assistance.

## Supplementary Figures

**Fig. S1.**
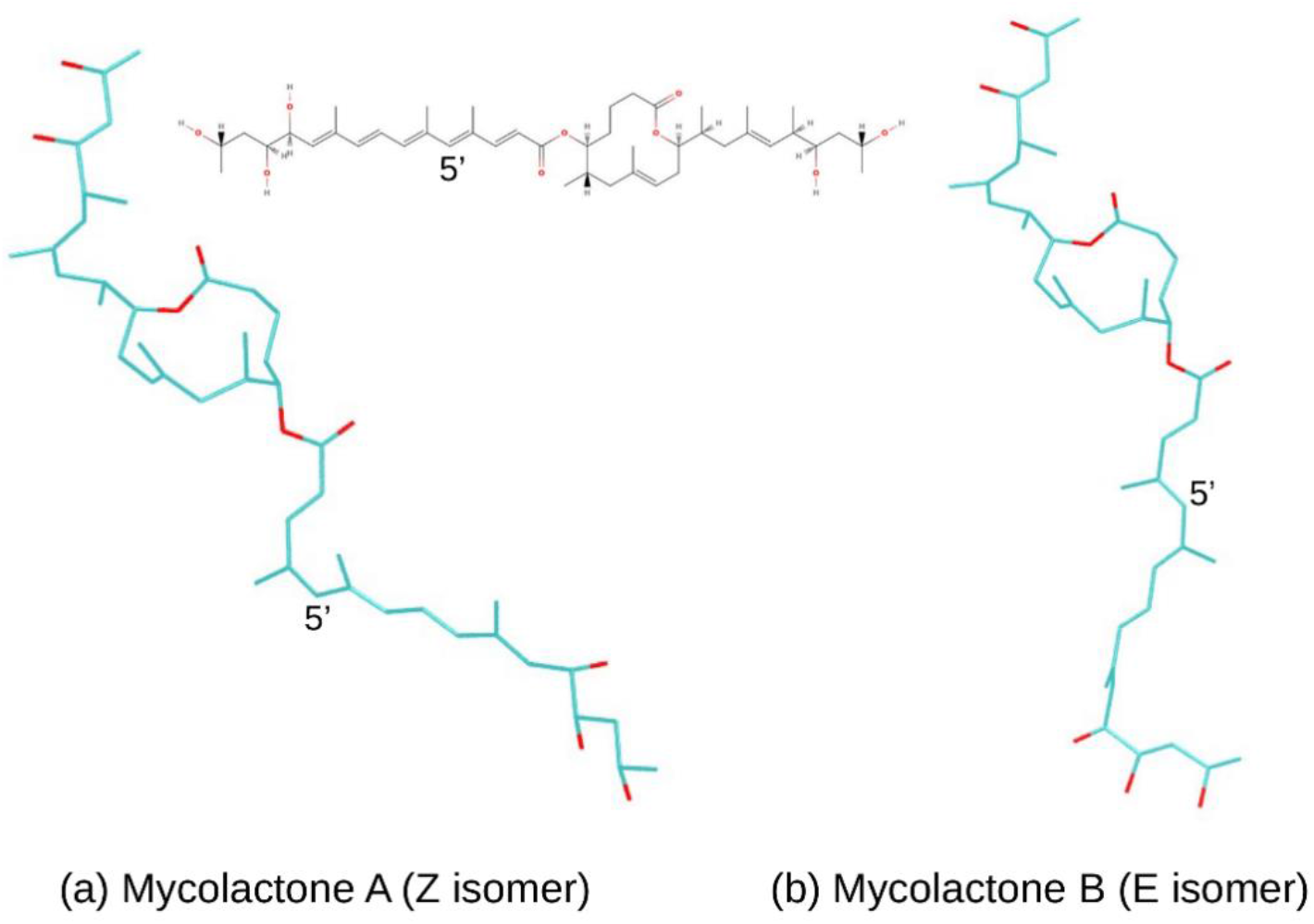
Conformations of mycolactone isomers.

**Fig. S2.**
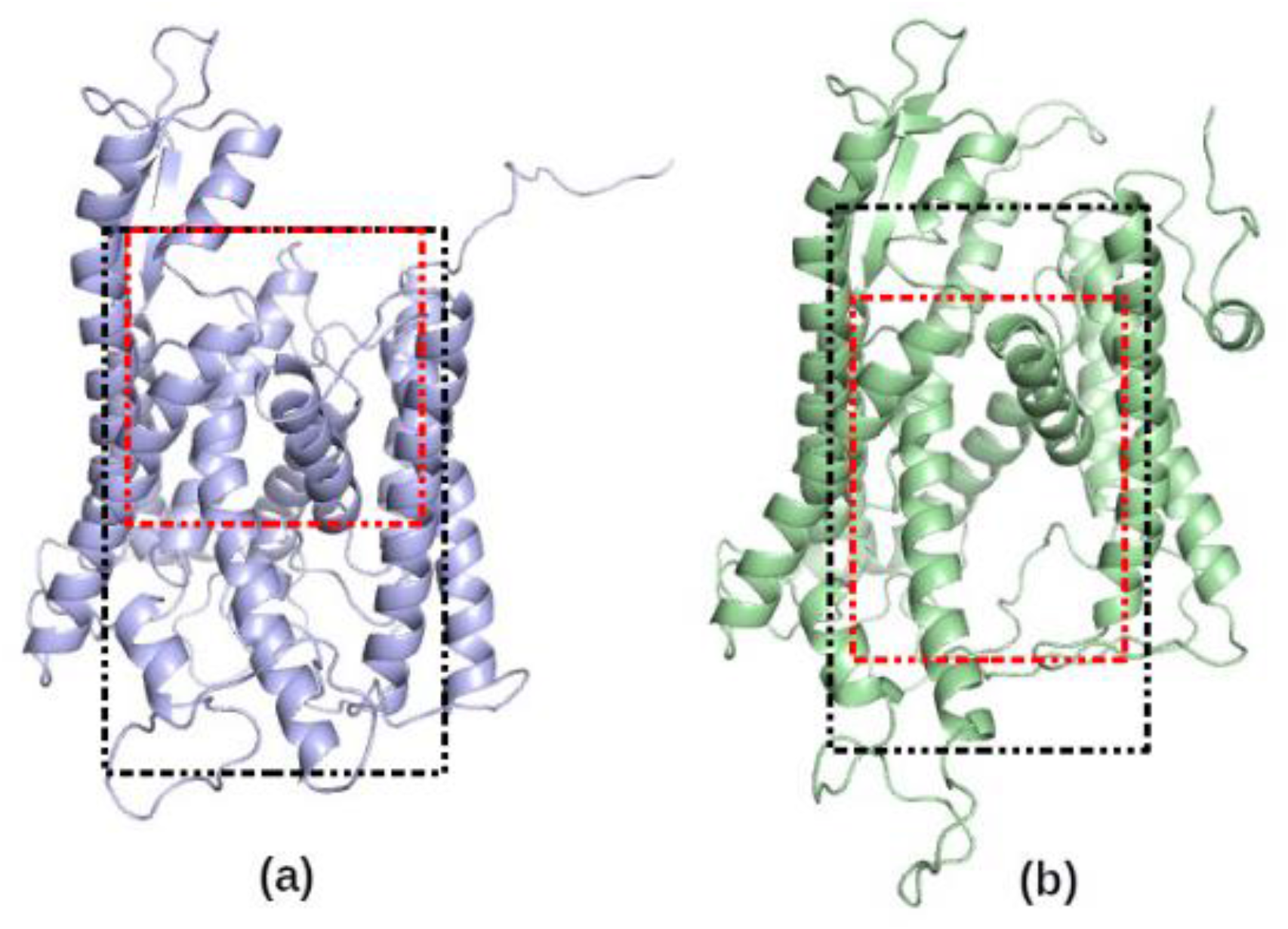
Docking regions in and around human Sec61α for (a) *idle* and *inhibited* states, and (b) for the *open* state. The black-dashed boxes represent the grid box (100 Å × 100 Å × 126 Å) used for the first stage of docking and the red-dashed boxes show the grid box used in the second (finer) docking stage (90 Å × 80 Å × 80 Å), (80 Å × 90 Å × 80 Å) and (80 Å × 80 Å × 90 Å) for *idle, intermediate* and *open* state, respectively. The conformations in (a) and (b) are homology models of *idle* and *open* states, respectively.

